# Sensitivity of the paired motor unit analysis for estimation of motoneuron excitability to commonly used constraints and filters

**DOI:** 10.1101/732982

**Authors:** Altamash Hassan, Christopher K. Thompson, Francesco Negro, Mark Cummings, Randy Powers, CJ Heckman, Jules Dewald, Laura Miller McPherson

**Author notes:** **Contact Information of Corresponding Author: (I can’t use Wash U in publications until I officially resign from FIU in Sept)** Laura McPherson, Department of Physical Therapy, Nicole Wertheim College of Nursing and Health Sciences, Florida International University, 11200 SW 8^th^ Street, AHC3-424, Miami, FL 33199.

## Abstract

The nervous system has a tremendous ability to modify motoneuron excitability according to task demands through neuromodulatory synaptic input to motoneurons. Neuromodulatory inputs adjust the response of the motoneuron to excitatory and inhibitory ionotropic input and can facilitate the induction of persistent inward currents (PICs). PICs amplify and prolong the motoneuron response to synaptic inputs, and PIC impairment may play a major role in motor deficits observed in pathological conditions. Noninvasive estimation of the magnitude of neuromodulatory input and persistent inward currents in human motoneurons is achieved through a paired motor unit analysis (ΔF) that quantifies hysteresis in the firing rates at motor unit recruitment and derecruitment. While the ΔF technique is commonly used for estimating motoneuron excitability, computational parameters used for the technique vary across studies. In the present study, we assessed the sensitivity of the ΔF technique to several criteria commonly used in selecting motor unit pairs for analysis, as well as to methods used for smoothing the instantaneous motor unit firing rates. Using HD-sEMG and motor unit decomposition we obtained 5,409 motor unit pairs from the triceps brachii of ten healthy individuals during submaximal triangle contractions. The mean (SD) ΔF was 4.9 (1.08) pps, consistent with previous work using intramuscular recordings. There was an exponential plateau relationship between ΔF and the recruitment time difference between the motor unit pairs, with the plateau occurring at approximately 1 s. There was an exponential decay relationship between ΔF and the derecruitment time difference between the motor unit pairs, with the decay stabilizing at approximately 1.5 s. We found that reducing or removing the minimum threshold for the correlation of the rate-rate slope for the two units did not affect ΔF values or variance. Additionally, we found that removing motor unit pairs in which the control unit was saturated had no significant effect on ΔF. Smoothing filter selection had no substantial effect on ΔF values and ΔF variance; however, the length and type of smoothing filter affected the minimum recruitment and derecruitment time differences. Our results facilitate interpretation of findings from studies that implement the ΔF approach but use different computational parameters.

## Introduction

Initial investigations of motoneuron firing patterns proposed that the output of a motoneuron is linear to the excitatory and inhibitory synaptic inputs the motoneuron receives. However, recent studies have shown that this relationship is non-linear due to the influence of neuromodulatory synaptic inputs. Serotonin (5-HT) and norepinephrine (NE) are robust monoaminergic neuromodulators that act through G-protein coupled receptors to dramatically change motoneuron excitability by adjusting the response of the motoneuron to excitatory and inhibitory ionotropic input (Heckman and Enoka 2012). These monoamines have a prominent effect on motoneuron dendrites by activating persistent inward currents (PICs), comprised of slow L-type Ca+ currents and fast persistent Na+ currents, which evoke a sustained depolarization in the cell (Hounsgaard, Hultborn et al. 1984, Bennett, Hultborn et al. 1998). This depolarization leads to amplified and prolonged responses in motoneuron output in relation to excitatory synaptic inputs, creating the distinctive firing patterns we see in motoneurons.

There is a small but growing body of recent work in humans that is beginning to reveal the importance that PICs have in both typical and pathological motor control. In the intact nervous system, the influence of PICs likely varies throughout the body and may be crucial in the control of muscles with different functions. For example, because the prolonged motoneuron output elicited by PICs is advantageous for muscles that must be activated for extended periods, postural and anti-gravity muscles are likely to have larger PICs than muscles specialized for fine motor control (Binder and Powers 2001, Heckman and Enoka 2012, Wilson, Thompson et al. 2015). Additionally, abnormal neuromodulatory synaptic input and/or PICs may underlie motor deficits seen in pathological states. In individuals with chronic spinal cord injury, uncontrolled muscle spasms and hyperactive reflexes have been linked to PICs elicited by constitutively active serotonin receptors (Gorassini, Knash et al. 2004, Li, Gorassini et al. 2004, Murray, Nakae et al. 2010, Murray, Stephens et al. 2011). In individuals with chronic stroke, increased monoaminergic drive and PICs may be partially responsible for the hyperactive stretch reflexes and the upper extremity flexion synergy in (McPherson, Ellis et al. 2008, McPherson, Ellis et al. 2018, McPherson, McPherson et al. 2018). Weakness associated with sepsis may be related to impaired PICs, as serotonin agonist-induced PICs have also been shown to ameliorate motor neuron firing deficits in a preclinical model of sepsis (Nardelli, Powers et al. 2017). This work in pathological populations emphasizes the role that neuromodulatory inputs and PICs play in the control of movement and the importance of their study. Nonetheless, much is still unknown and further study of neuromodulatory inputs and PICs is necessary.

Although PICs cannot be directly measured from human motoneurons, experimental techniques have been developed to estimate the size of PICs in humans via motor unit recordings. Currently, the standard method for estimating PIC amplitude (thus allowing for inference of neuromodulatory synaptic input) is the ΔF technique developed by Gorassini and colleagues (Gorassini, Yang et al. 2002). With this technique, PIC amplitude is estimated by quantifying motor unit recruitment/de-recruitment hysteresis using pairs of motor units firing during slow linear “triangle” contractions. The ΔF metric has been validated through both animal and simulation work (Powers, Nardelli et al. 2008, Powers and Heckman 2015) and has shown sensitivity to increased monoaminergic drive in humans given amphetamines (Udina, D’Amico et al. 2010).

Conventionally, the ΔF technique requires that motor unit pairs meet certain criteria based on assumptions related to the underlying physiology. For example, the difference in recruitment time between the control and test unit must be long enough to ensure that the PIC in the control unit is fully active before test unit recruitment. Additionally, the lower threshold (control) and higher threshold (test) units must have sufficient correlation of their firing rates, as the firing rate of the control unit is used as an approximation of the ionotropic excitatory synaptic input to the test unit. Also, firing rate saturation in the control unit may bias the ΔF calculation and is often controlled for in these analyses. Despite the use of these standard criteria, the specific parameter values for each criterion vary across studies. Further, there are differences in computational factors across studies, such as the type of smoothing filter used on the motor unit spike times.

The purpose of the present study is to determine the sensitivity of the ΔF technique to differences in 1) minimum recruitment and derecruitment time difference; 2) minimum rate-rate slope correlation; 3) control unit firing rate modulation; and 4) filter selection. Such a robust sensitivity analysis is now possible as we can obtain spike trains from large populations of motor units using the high-density surface EMG (HD-sEMG) decomposition approaches (Holobar and Zazula 2007, Chen and Zhou 2015, Negro, Muceli et al. 2016). Previous work with ΔF has largely used intramuscular recordings and has therefore been limited by the number of motor units that can be feasibly recorded. Here, we present motor unit data obtained using convolutive blind source separation of HD-sEMG signals (Negro, Muceli et al. 2016) recorded from the human triceps.

## Methods

### Participants

Ten adults (3 female, 7 male) ranging in age from 22 to 31 (mean ± SD age: 26.2 ± 2.4) completed the study. For inclusion in this study all participants were required to have: (1) no known neurological injury or disease, (2) no muscular impairment of upper extremity motor function, and (3) no significant visual or auditory impairments. All participants provided written informed consent prior to participation in this experiment which was approved by the Institutional Review Board of Northwestern University.

### Experimental Apparatus

Participants were seated in a Biodex experimental chair (Biodex Medical Systems, Shirley, NY) and secured with shoulder and waist straps to minimize trunk movement. In order to measure isometric elbow torques, the participant’s dominant forearm was placed in a fiberglass cast and rigidly fixed to a six degrees-of-freedom load cell (JR3, Inc., Woodland, CA). The arm was positioned at a shoulder abduction (SABD) angle of 75° and an elbow flexion angle of 90°. The fingers were secured to a custom hand piece at 0° wrist and finger (metacarpophalangeal) flexion/extension (Miller, Thompson et al. 2014) (Figure 1).

**Figure 1:**
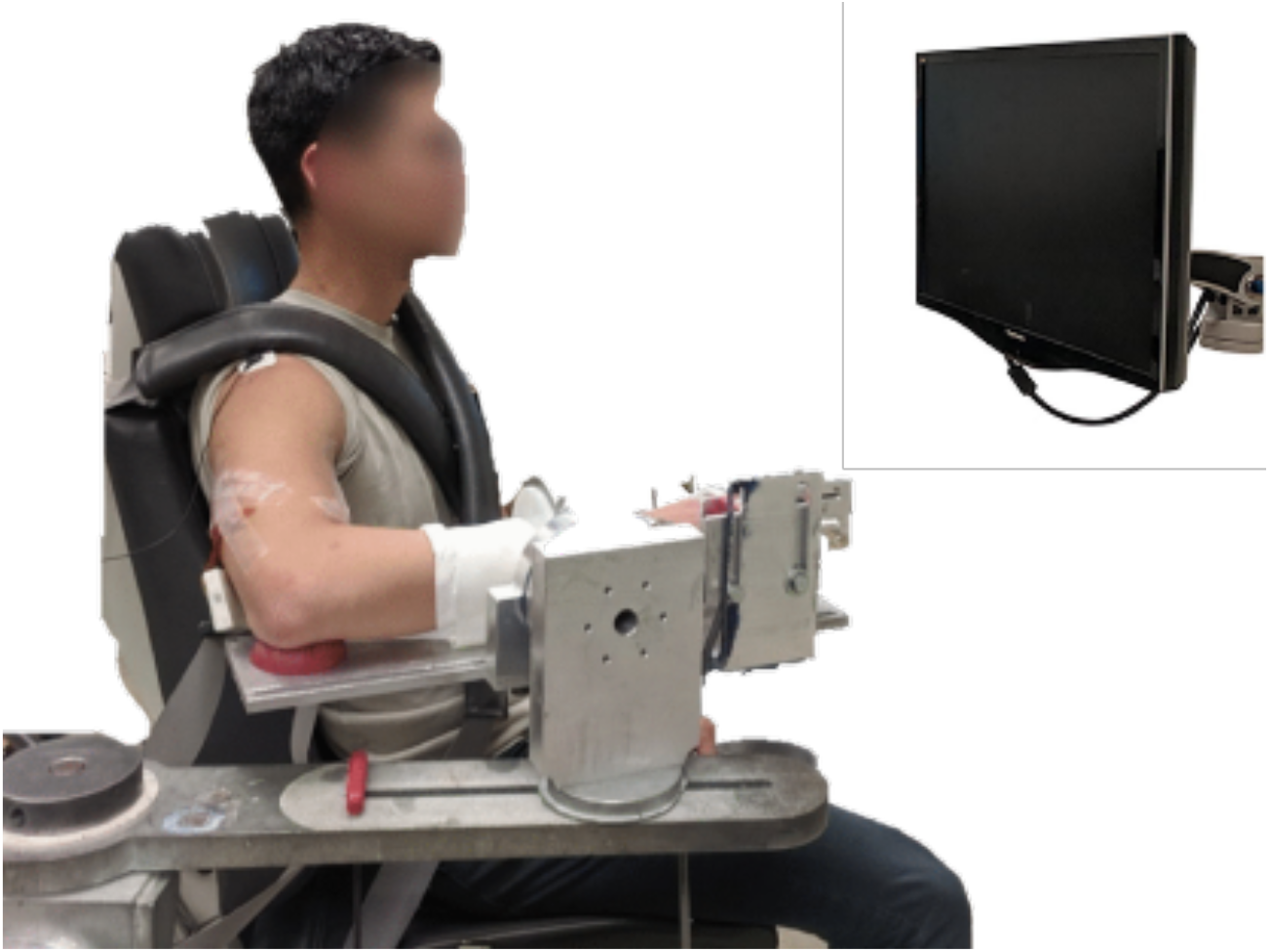
Isometric joint torque recording device with high-density surface EMG grids on the biceps brachii and triceps brachii.

Forces and torques measured at the forearm-load cell interface were recorded at 1024 Hz and converted into elbow flexion and extension torques through a Jacobian based algorithm, utilizing limb segment lengths and joint angles, implemented by custom MATLAB software (The MathWorks).

Multi-channel surface EMG recordings were collected in single differential mode from the lateral head of the triceps brachii using a grid of 64 electrodes with 8mm inter-electrode distance (GR08MM1305, OT Bioelettronica, Inc.) (Figure 1). The signals were amplified (x150), band-pass filtered (10-500Hz) and sampled at 2048 Hz (Quattrocento, OT Bioelettronica, Turin, IT). The EMG recordings and the force/torque recordings were collected on separate computers; a 1 second TTL pulse was transmitted to both computers for use as a marker, and each trial was temporally synced offline using a cross-correlation of the TTL pulses.

### Protocol

First, participants were asked to generate maximum voluntary torques (MVTs) in the direction of elbow extension. Real time visual feedback of torque performance was provided on a computer monitor. Trials were repeated until three trials in which the peak torque was within 10% of each other were collected. If the final trial had the highest peak torque, a subsequent trial was collected. Participants were provided with enthusiastic vocal encouragement during MVT trials and were given adequate rest breaks between trials to prevent fatigue.

Experimental trials entailed the generation of triangular isometric torque ramps using real-time visual feedback of elbow flexion/extension torque. Participants were instructed to gradually increase their elbow extension torque to ~20% MVT over 10 seconds and then gradually decrease their torque back to 0% MVT over the subsequent 10 seconds. Each trial consisted of either two or three ramps in succession, with ten seconds of rest between ramps and five seconds of rest at the beginning and end of each trial. Participants were given several practice trials to become comfortable with the task, followed by five to six experimental trials that were used for subsequent analysis. Torque traces were visually inspected and trials with large deviations from the desired time-torque profile were discarded.

### Motor unit decomposition and selection

All surface EMG channels were visually inspected and those with substantial artifacts or noise were removed (typically zero to five channels were removed per trial). The remaining surface EMG channels were decomposed into motor unit spike trains based on convolutive blind source separation (Negro, Muceli et al. 2016) and successive sparse deflation improvements (Martinez-Valdes, Negro et al. 2017). The silhouette threshold for decomposition was set to 0.85. However, even with this high threshold of decomposition accuracy, the blind source separation algorithm may still extract some solutions which do not relate to physiological motor unit firing patterns. To address these errors, we supplemented the automatic decomposition with visual inspection of the motor unit firings and iterative improvement by experienced investigators with the use of a custom-made graphical user interface. This approach has been previously applied (Boccia, Martinez-Valdes et al. 2019) and provides a high degree of accuracy in the estimated discharge patterns.

Motor unit firing times were obtained from the decomposed spike trains. Motor unit firing rates were calculated as the reciprocal of the time between consecutive motor unit firings, or inter spike interval (ISI).

While the automatic blind source separation does not produce any duplicate motor units, the visual inspection and iterative improvement process occasionally leads to duplicate units due to the separation of merged motor units or removal of erroneous firing times. To ensure that duplicate motor units were detected and eliminated, the spike times of motor units within the same muscle and trial were cross correlated. If any motor unit pairs shared more than a 50% overlap in spike times, the motor unit with the higher covariance value was removed.

### Paired Motor Unit Analysis

ΔF is a paired motor unit analysis that quantifies the effects of PICs on motoneuron firing patterns by measuring the discharge hysteresis of a higher threshold motor unit (test) with respect to the firing rate of a lower threshold motor unit (control) (Gorassini, Yang et al. 2002). Specifically, ΔF of the test motor unit is calculated as the difference in the firing rate of the control motor unit between the time of recruitment and derecruitment of the test motor unit. Figure 2 illustrates this method of analysis.

**Figure 2:**
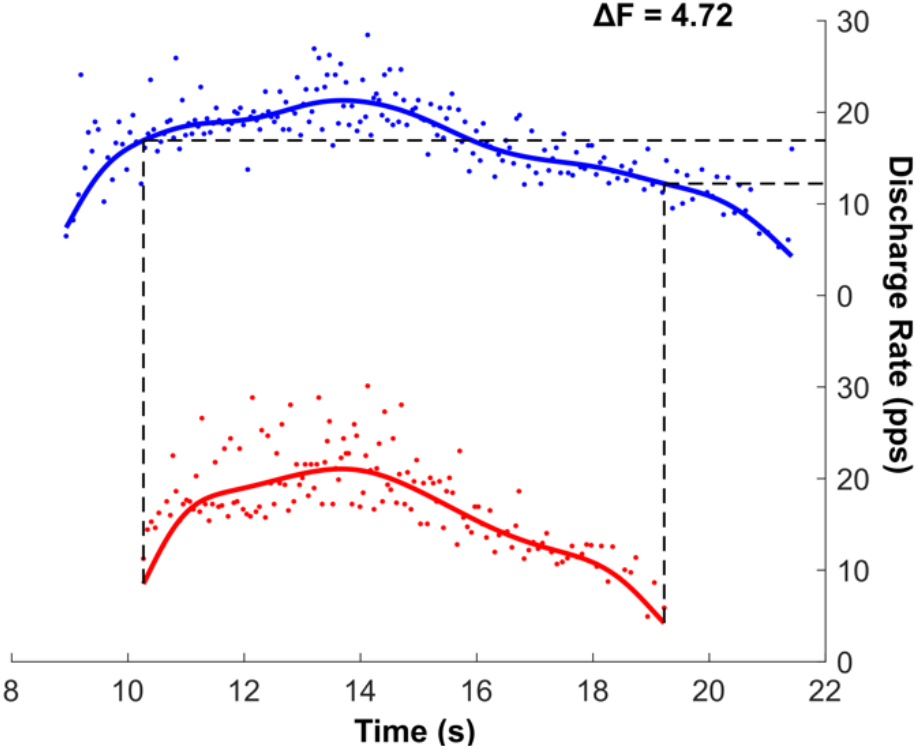
An example of the F technique where the change in firing rate of the control unit (blue) is measured at the recruitment and derecruitment of the test unit (red) taken from the triceps brachii of a young control participant. The solid blue and red lines

The ΔF technique first considers every combination of motor unit pairs in which the lower threshold control unit fires through both recruitment and derecruitment of the higher threshold test unit. The test unit must fire for at least 2 s to ensure the PIC can be fully activated (Stephenson and Maluf 2011). Then, the motor unit pairs must meet additional criteria to be appropriate for further analysis.

Criteria commonly used for the ΔF technique are the following: (1) a minimum threshold for the time difference between recruitment of the motor unit pairs, (2) a minimum rate-rate slope (reflecting sufficient shared synaptic input), and (3) sufficient rate modulation in the control unit. Here, we assess the sensitivity of the ΔF calculation to various parameter values of these criteria.

#### Recruitment time difference

The criterion of a minimum recruitment time difference between the control and test motor units is based on the idea that the PIC in the control unit must be fully activated prior to the recruitment of the test unit. Early work required a minimum of 2 s between recruitment of the control and test unit based on initial literature showing the PIC can take up to 2 s to fully activate (Hounsgaard and Kiehn 1989, Bennett, Li et al. 2001, Li, Gorassini et al. 2004). However, simulation work by Powers and Heckman (Powers and Heckman 2015) has suggested that the effect of recruitment time difference on ΔF and its variance across motor unit pairs diminishes greatly after 0.5 s.

#### Derecruitment time difference

The minimum derecruitment time difference between the control and test units may also have a substantial effect on ΔF, as PIC inactivation in the control unit may affect the ΔF calculation. Previous work has not investigated the effect of derecruitment time difference on the ΔF analysis; however, deactivation of the PIC in the control unit very close in time to deactivation of the test unit could lead to overestimation of PICs.

#### Rate-rate slopes

The ΔF calculation relies on the assumption that both the control and test unit share substantial synaptic input as quantified using the correlation in the rate-rate slopes. A consistent limit for rate-rate slope correlation has not been established. The initial threshold of r^2^ ≥ 0.7 used by Gorrassini and colleagues (Bennett, Li et al. 2001, Gorassini, Yang et al. 2002) has been used extensively (Udina, D’Amico et al. 2010, Stephenson and Maluf 2011, Wilson, Thompson et al. 2015). However, other work has used more lenient limits on rate-rate slope correlation including r^2^ ≥ 0.6 (Mottram, Suresh et al. 2009) and r^2^ ≥ 0.5 (Powers, Nardelli et al. 2008), and r^2^ ≥ 0.3 (Zijdewind, Bakels et al. 2014). We investigated the effect of 8 different rate-rate slope correlation minima (r^2^ > 0, 0.25, 0.5, 0.7, 0.75, 0.8, 0.85, 0.9) on the ΔF calculation.

#### Rate modulation of the control unit

If the firing rate of the control unit does not reflect the net ionotropic synaptic drive (e.g., due to decreased rate modulation of that unit), then the PIC amplitude using the ΔF method could be underestimated. Rate modulation in the control unit is here defined as the range of firing rates of the control unit during the time the test unit is active. Previous work (Stephenson and Maluf 2011, Wilson, Thompson et al. 2015) has excluded motor unit pairs in which the rate modulation of the control unit is within 0.5 pps of the calculated ΔF, to ensure rate saturation of the control unit is not limiting ΔF. Here we evaluated the effect of removing motor unit pairs which showed control unit saturation on the ΔF calculation.

#### Filter selection

Variation in computational methods used to prepare motor unit firing patterns for the ΔF analysis may affect the results. Gorassini and colleagues’ original implementation of the ΔF method fit a 5^th^-order polynomial to the instantaneous firing rates (Gorassini, Yang et al. 2002) while previous motor unit work has filtered instantaneous firing rates using a Hanning window (De Luca and Erim 1994, de Luca, Foley et al. 1996, De Luca and Contessa 2012) or a Gaussian window (Powers, Nardelli et al. 2008). These smoothing methods have different effects on the firing patterns, particularly as a result of edge effects at motor unit recruitment and derecruitment. The shape of Hanning and Gaussian filters may produce sharp downward edges, while the 5^th^-order polynomial is more sensitive to doublets and errors, which may disproportionally skew the polynomial fit at recruitment and derecruitment. Here we compare ΔF values calculated using a 1s Hanning window, a 2 s Hanning window, a 2 s Gaussian window, and a 5^th^ order polynomial to smooth instantaneous firing rates.

#### Approach to sensitivity analysis

We first examined the effect of recruitment and derecruitment time difference on ΔF using 3 different rate-rate correlation thresholds and a 2s Hanning window, which has been used extensively by our group and many other investigators(De Luca and Erim 1994, De Luca and Contessa 2012), to smooth motor unit firing patterns. The minimum time differences obtained from the recruitment and derecruitment time difference were used for the subsequent rate-rate correlation analysis. The results from the recruitment and derecruitment time difference analysis and the rate-rate correlation analysis were used for both the analyses of control unit saturation and filter selection.

## Results

In total, 1576 motor unit spike trains were decomposed from the triceps brachii of 10 participants. Each participant completed at least 8 isometric elbow extension triangle contractions with an average yield of 10.4 ± 4.3 motor units per trial. We considered 5,409 motor unit pairs for the ΔF analysis. A small number of these pairs (106) were excluded because the test unit was active for less than 2 s.

### Relation of ΔF values to recruitment and derecruitment time difference

Figure 3 a-c shows the relationship between ΔF values and the time difference between control and test unit recruitment with three different thresholds for rate-rate correlation (r^2^ > 0.5, 0.7, 0.9). With all three rate-rate correlation thresholds, the ΔF values demonstrated an exponential plateau behavior, rapidly increasing along with recruitment time difference values before plateauing. To approximate the minimumrecruitment time difference at which ΔF values no longer varied, we fit the data using an exponential plateau function and identified where the exponential fit had grown 3 half-lives, reaching 87.5% of its asymptotic value. This resulted in a recruitment time difference cutoff of 0.92 s, 0.95 s, and 0.91 s for the motor unit pairs with r^2^ > 0.5, 0.7, and 0.9, respectively.

**Figure 3:**
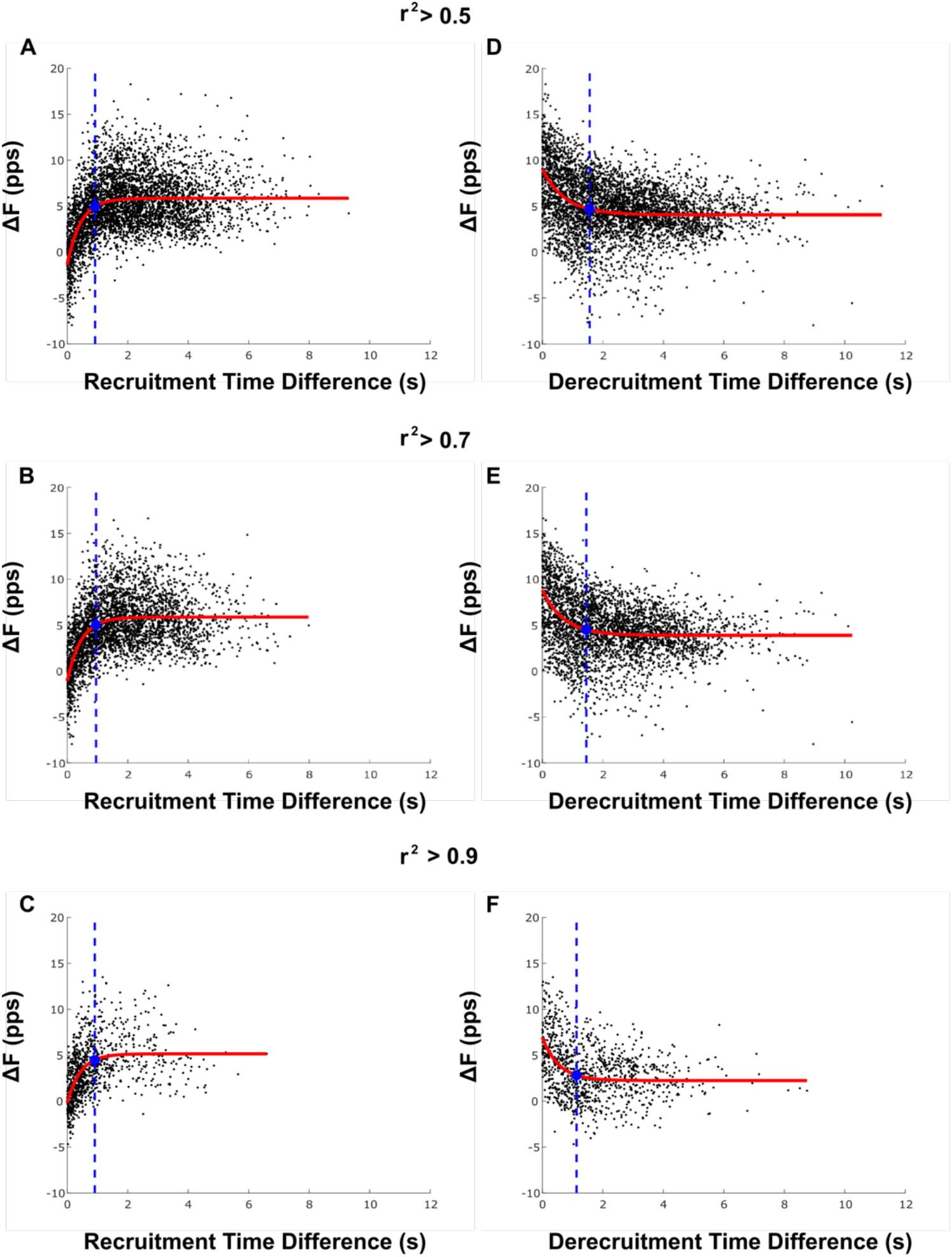
The relationship between recruitment time difference and ΔF at 3 different thresholds for rate-rate correlation (A,B,C). The relationship between derecruitment time difference and ΔF at 3 different thresholds for rate-rate correlation (D,E,F). Red line denotes exponential plateau fit. Blue filled circles denote the 87.5% of the asymptote model, and blue dotted line denotes the minimum recruitment/derecruitment time used for further analyses.

Figure 3 d-f shows the relationship between ΔF values and the time difference between test and control unit derecruitment, with 3 different thresholds for rate-rate correlation (r^2^ > 0.5, 0.7, 0.9). A decaying exponential plateau function was used to fit the data, and the minimum derecruitment time difference was determined as the point where the exponential fit had decayed to 87.5% of its asymptotic value. The minimum derecruitment time difference was 1.56 s, 1.45 s, and 1.13 s for the motor unit pairs with r^2^ > 0.5, 0.7, and 0.9, respectively.

Based on these results, we restricted our following analyses to motor unit pairs with at least 1 s difference between control and test unit recruitment times and 1.5 s difference between test and control unit derecruitment times, which yielded a mean +/- SD of 304.1 ± 178.4 motor unit pairs per participant.

### Dependence of ΔF on rate-rate slope correlation

The average number and percentage of motor unit pairs with rate-rate slope correlation values above each threshold (r^2^ > 0, 0.25, 0.5, 0.7, 0.75, 0.8, 0.85, 0.9) are shown in Table I. The percentage of retained motor unit pairs decreased dramatically when the r^2^ threshold increased beyond 0.5, dropping from 76.1% with r^2^ > 0.5 to 49.8% with r^2^ > 0.7 to only 6.5% with r^2^ > 0.9.

**Table I:** Number and percentage of motor unit pairs at various rate-rate slope correlation thresholds (group mean ±SD)

| Minimum $r^2$ | Number of pairs | % of pairs |
| --- | --- | --- |
| 0.00 | 304.1 ± 178.4 | 100 ± 0.00 |
| 0.25 | 276.6 ± 164.8 | 90.7 ± 0.07 |
| 0.50 | 232.0 ± 146.7 | 76.1 ± 0.14 |
| 0.70 | 160.2 ± 124.2 | 49.8 ± 0.21 |
| 0.75 | 133.1 ± 110.8 | 40.9 ± 0.21 |
| 0.80 | 98.8 ± 83.9 | 30.5 ± 0.16 |
| 0.85 | 55.8 ± 49.7 | 17.6 ± 0.10 |
| 0.90 | 19.7 ± 17.5 | 6.5 ± 0.04 |

Figure 4 A shows the relationship between group mean ΔF value and each rate-rate slope correlation threshold. Figure 4 B displays the group mean individual participant variance in ΔF across the different thresholds for rate-rate slope correlation. At the majority of the rate-rate slope correlation thresholds (below 0.85), both the group mean ΔF values and group mean individual participant variance were remained consistent. The group mean ΔF value of ~5 pps is comparable to results from previous work that calculated ΔF in the triceps brachii using motor unit data obtained using intramuscular EMG decomposition (Wilson, Thompson et al. 2015). At the strictest r^2^ thresholds (r^2^ > 0.85 and r^2^ > 0.9), the group mean ΔF value decreased, and the group mean individual participant variance increased.

**Figure 4:**
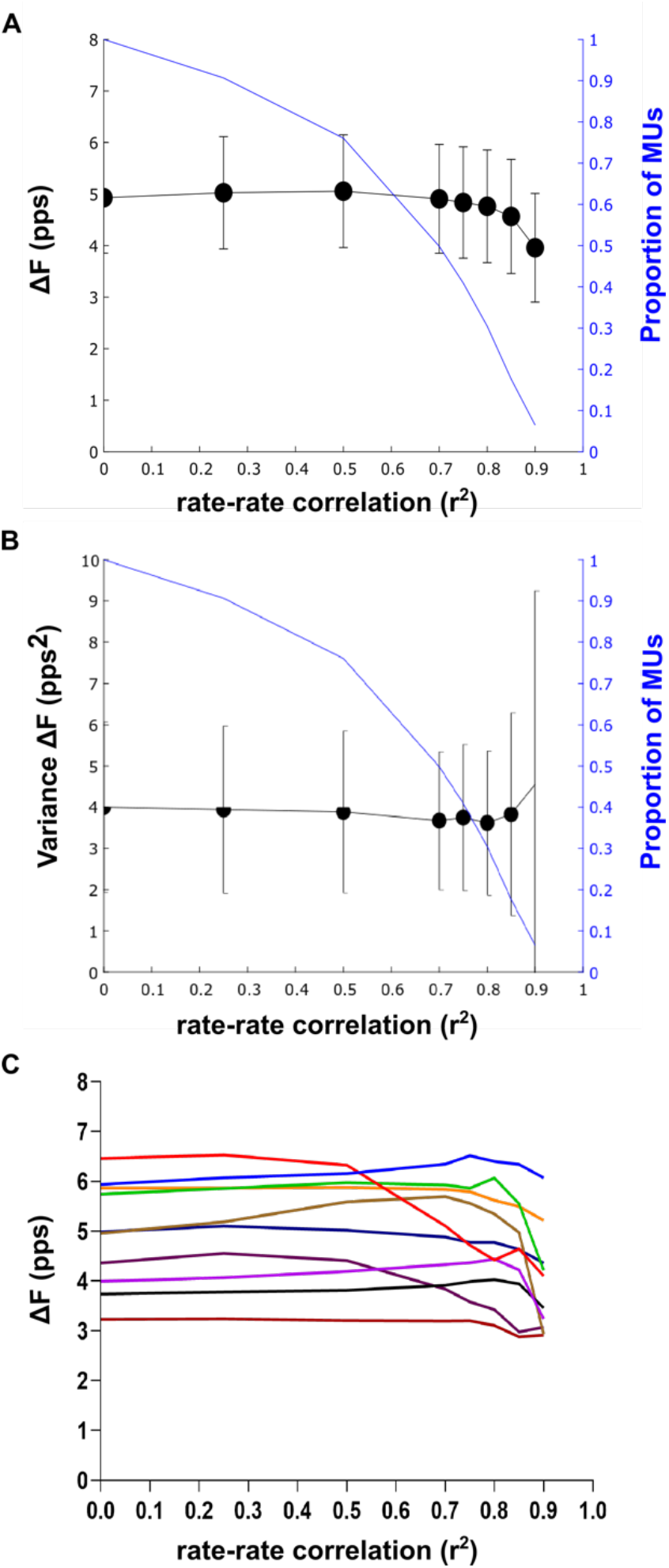
The group mean ± SD ΔF values (A) and group mean individual participant variance in ΔF (B) across three different thresholds for rate-rate correlation. The relationship between ΔF and minimum rate-rate correlation for each of the 10 participants is shown in part C.

Figure 4 C shows the relationship between ΔF value and minimum rate-rate slope correlations for each participant. ΔF values for all participants were relatively stable for r^2^ thresholds of up to 0.5. For three participants, ΔF decreased markedly with higher r^2^ thresholds whereas values for the other participants fluctuated to a lesser degree.

Similar ΔF values and variance were obtained when using no rate-slope correlation threshold as were obtained when using the traditional rate-rate correlation restrictions. Based on this stability, as well as the higher number of motor unit pairs afforded by removing this restriction, we removed the minimum rate-rate correlation restriction in our following analyses.

### Dependence of ΔF on control unit firing rate modulation

The maximum ΔF value of a motor unit pair is limited by the amount of rate modulation in the control unit during test unit firing. In order to avoid underestimation of ΔF due to insufficient rate modulation in the control unit, previous studies have removed motor unit pairs in in which the ΔF value was within 0.5 pps of the control unit rate modulation (Stephenson and Maluf 2011, Wilson, Thompson et al. 2015). Figure 5 A shows the relationship between ΔF and the firing rate modulation of the control unit; motor unit pairs which fit the criteria for control unit saturation are shown in blue. Figure 5 B shows the group mean ΔF before and after the removal of motor unit pairs that exhibited possible saturation. The group mean ΔF was 4.9 ± 1.08 pps before removal of pairs which exhibited control unit saturation, and the group mean ΔF was 4.7 ± 0.96 pps following the removal of those pairs. There was no significant change in subject mean ΔF after removal of pairs which fit the saturation criterion (P = 0.17). Figure 6 C shows the mean ΔF per subject before and after the removal of motor unit pairs that exhibited possible saturation.

**Figure 5.**
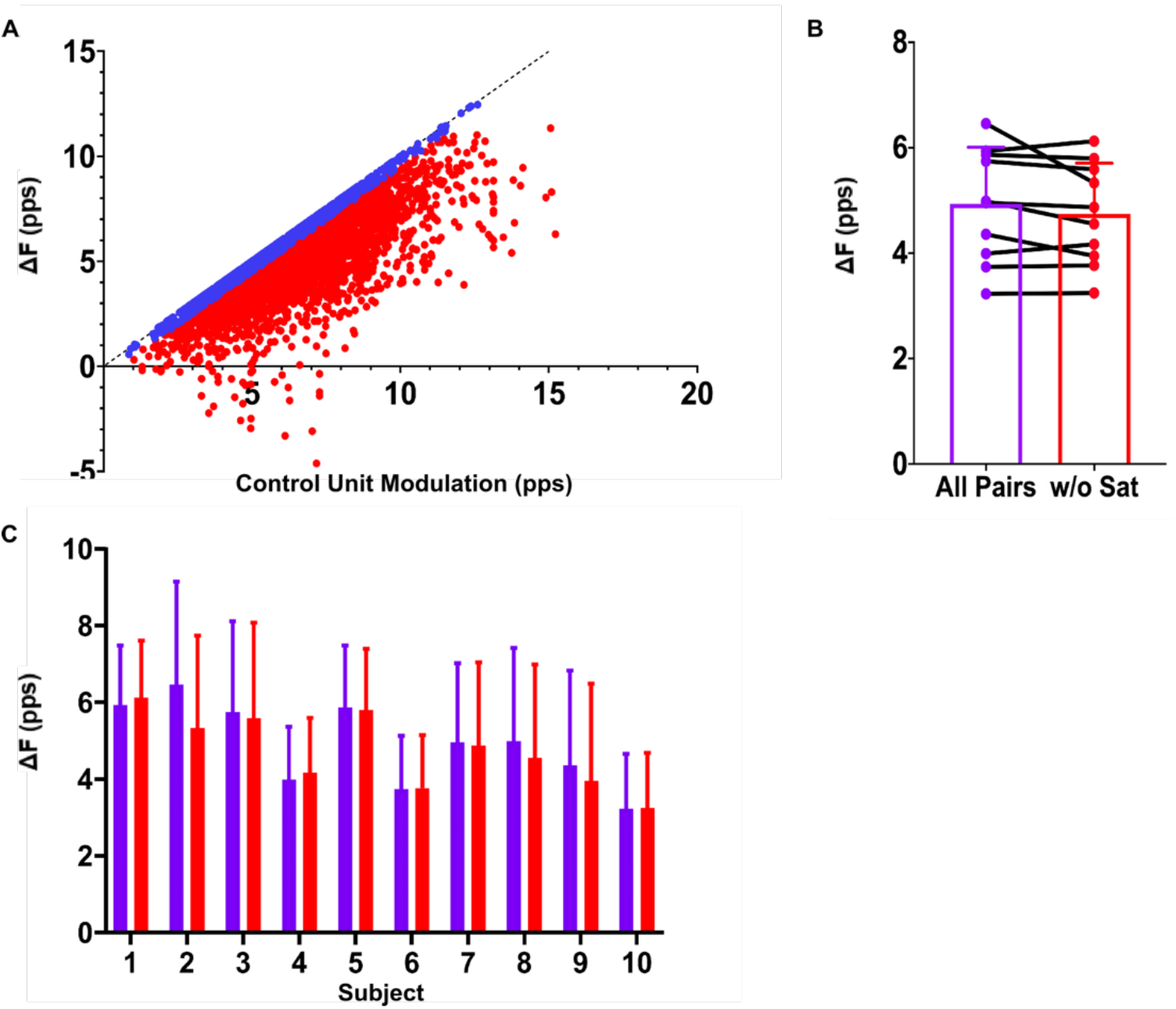
A: the relationship between ΔF and control unit firing rate modulation with motor unit pairs matching the criteria for control unit saturation shown in blue. B: the group mean ± SD and individual participant ΔF values before and after the removal of motor unit pairs displaying control unit saturation. C: the mean ± SD ΔF per subject before (purple) and after (red) the removal of motor unit pairs with control unit saturation.

**Figure 6.**
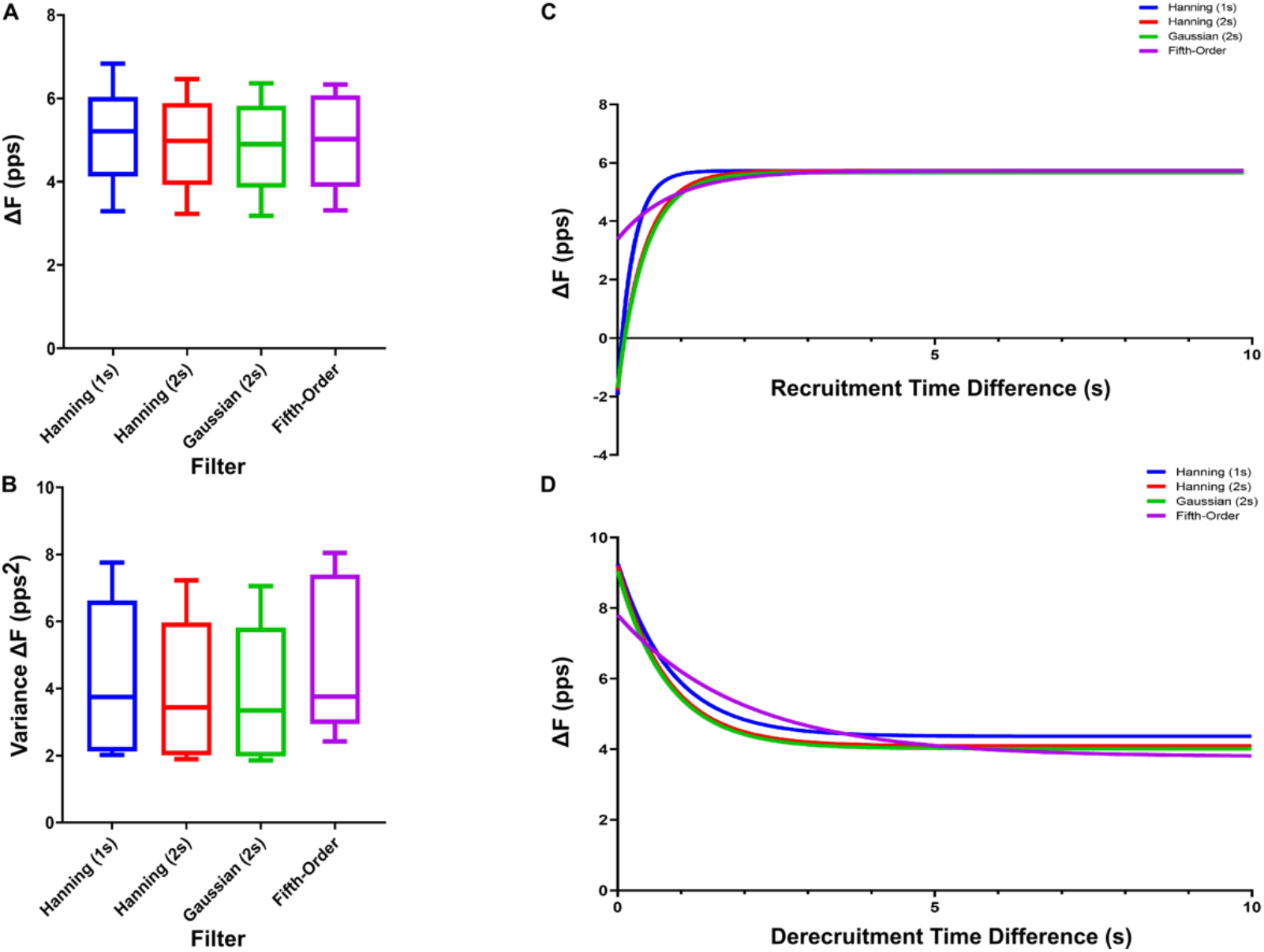
Subject mean ΔF (A) and mean subject variance (B) plotted across four filter types. An exponential plateau function showing the relationship between ΔF and recruitment time difference (C) and derecruitment time difference (D) for the same four filter types.

### Effect of filter selection on ΔF results

The ΔF technique relies on filtering of instantaneous motor unit firing rates to provide smoothed continuous firing rates. Figure 6 A shows the change in subject mean ΔF across 4 different filter methods. The group mean ΔF was 5.1 ± 1.12 pps for the 1 s Hanning window, 4.9 ± 1.08 pps for the 2 second Hanning window, 4.9 ± 1.06 pps for the 2 second Gaussian window, and 5.0 ± 1.12 pps for the fifth-order polynomial fit. While a one-way ANOVA reveals a significant effect of filter on subject mean ΔF value (P < 0.0001), the amplitude of the difference between filter types is minute. Figure 6 B shows the relationship between the mean subject variance and filter type. The mean subject variance is 4.1 ± 2.29 pps^2^ for the 1 s Hanning window, 4.0 ± 2.06 pps^2^ for the 2 s Hanning window, 3.9 ± 2.01 pps^2^ for the 2 s Gaussian window, and 4.9 ± 2.31 pps^2^ for the fifth-order polynomial fit. Similar to the subject mean ΔF values, a one-way ANOVA reveals a significant effect of filter type on subject mean variance (P < 0.0001).

The edge effects of the different smoothing methods may affect the relationship between ΔF values and recruitment and derecruitment time difference. To investigate these effects, an exponential plateau function was used to fit the relationship between recruitment time difference and ΔF, for all 4 smoothing methods. Figure 6 C shows the modeled relationships between ΔF and recruitment time difference for each filter type. The recruitment time difference where the exponential fit reached 87.5% of its asymptotic value, was shorter for the data smoothed with a 1 s Hanning window, 0.50 s, than the 2 s Hanning and Gaussian windows, 0.87 s and 0.91 s respectively. For the data smoothed with a fifth-order polynomial, the time difference where the exponential fit reached 87.5% of its asymptotic value was much larger, 1.82s. However, the data smoothed with a fifth order polynomial was less sensitive to recruitment time difference with an exponential fit range of 2.36 pps across the range of observed recruitment time differences, compared to 7.70 pps, 7.53 pps, and 7.39 pps for the 1 s Hanning window, 2 s Hanning, and 2 s Gaussian window, respectively.

Figure 6 D shows the modeled relationships between ΔF and derecruitment time difference. The derecruitment time difference where the fit reached 87.5% of its asymptotic value, was similar for the data smoothed with a 1 s Hanning window, 2 s Hanning window, and 2 s Gaussian window, 1.78 s, 1.63 s, and 1.65 s respectively, but was much later for the data smoothed with the fifth-order polynomial, 4.09 s. The exponential fit range for derecruitment time difference is 4.90 pps for the 1 s Hanning window, 5.11 pps for the 2 s Hanning window, 5.04 pps for 2 s Gaussian, and 4.02 pps for fifth-order polynomial.

## Discussion

In this study we utilized HD-sEMG and motor unit decomposition to quantify the relationship between the ΔF technique and its commonly used criteria, as well as the smoothing methods applied to motor unit firing patterns. Our average ΔF values (4.9 ± 1.08 pps) are similar to those measured using intramuscular EMG (Wilson, Thompson et al. 2015). We confirmed and further quantified the relationship between ΔF and recruitment time difference, which has been previously investigated (Gorassini, Yang et al. 2002, Stephenson and Maluf 2011, Wilson, Thompson et al. 2015). Further, we found a relationship between ΔF and the derecruitment time difference between the test and control units. We found ΔF values and variance were mostly independent of rate-rate correlation criteria, and only affected if tight restrictions on rate-rate correlation are used. We found that ΔF values and variance were relatively independent of the method used to smooth the motor unit firing rates; however, the filter methods affect the necessary recruitment and derecruitment time spacing of motor unit pairs. Additionally, we saw no effect of removing possibly saturated motor unit pairs on ΔF values.

### Effect of recruitment and derecruitment time difference on ΔF

The ΔF technique requires that the PIC of the control unit be active for the duration of test unit firing, to ensure the control unit firing rate varies linearly in response to changes in net excitatory input. If the PIC in the control unit has not been fully activated before the recruitment of the test unit ΔF may be underestimated. Previous studies have controlled for this by discarding motor unit pairs with recruitment time differences below a certain minimum, however, these thresholds vary across studies from 0.5 to 2 s.

In alignment with previous work (Powers, Nardelli et al. 2008, Stephenson and Maluf 2011, Wilson, Thompson et al. 2015), we observed a reduction in ΔF values for motor unit pairs with closely recruited control and test units. Further, we found an exponential plateau relationship between ΔF and recruitment time difference (Figure 3). While previous work has modeled this relationship with linear (Wilson, Thompson et al. 2015) or quadratic (Stephenson and Maluf 2011) fits, increased number of motor unit pairs across a wider range of recruitment time differences in this study show a plateau in ΔF values as recruitment time difference increases. Based on this exponential plateau relationship, we found a minimum recruitment time difference of ~ 1 s. This time course is similar to that of PIC activation recorded from intracellular recordings in rat motoneurons (Bennett, Li et al. 2001).

Additionally, an exponential decay relationship was observed between ΔF and the derecruitment time difference of the test and control units. When the control unit is derecruited closely after the test unit, ΔF may be overestimated. The increased ΔF for motor unit pairs with low derecruitment time differences suggests a deceleration of control unit firing rate near derecruitment. One possible explanation for the effect of derecruitment time difference on ΔF is PIC inactivation near derecruitment may cause the rapid deceleration in motor unit firing rate. Additionally, the edge effects of filters used to smooth instantaneous firing rates may also cause a sharper deceleration near derecruitment. Results from this study provide evidence of an effect of derecruitment time difference on ΔF, which should be controlled for in future studies.

### Relation between ΔF and rate-rate slope correlation

As the ΔF technique uses the control unit as an estimate of excitatory synaptic drive to the test unit, previous studies only use motor unit pairs which have strong correlation in their firings rates. Previous work has commonly used rate-rate correlation thresholds of r^2^ > 0.5-0.7.

The present study found reducing or removing the minimum threshold for rate-rate slope correlation did not affect ΔF value or its variance. These results are consistent with findings from the decerebrate cat (Powers, Nardelli et al. 2008). As previously posited, one possible explanation for these results is that the ΔF calculation only measures the control unit firing at two points, test recruitment and derecruitment (Powers, Nardelli et al. 2008). Differences in modulation of the test and control unit that do not occur at test recruitment and derecruitment would affect the rate-rate slope correlation, but not the ΔF value, as long as the control unit is a sensitive indicator of synaptic input at the onset and offset of discharge of the test unit.

While ΔF values were stable across lower minimum correlation thresholds, our results suggest putting stricter limitations on firing rate correlation leads to a decrease in ΔF value and increase in variance. The increased variance is likely due to the reduced number of motor units available for these analyses, shown in Table I and Figure 4. Selection bias may play a role in the reduced ΔF values observed with higher rate-rate correlation threshold. Motor unit pairs with higher firing rate correlation are often recruited closely together, which can lead to reduced ΔF. Additionally, only test units with minimal PIC-induced firing rate nonlinearities would have sufficiently high correlation with control units which have fully activated PICs, limiting the selection to units with lower ΔF.

Relaxing the limits on rate-rate slope correlation may enable the calculation of ΔF values in pathological conditions that may alter motor unit firing rate correlation, such as muscle spasms in individuals with chronic spinal cord injury (Zijdewind, Bakels et al. 2014).

### Effect of control unit firing rate modulation on ΔF

Due to the nature of the ΔF calculation, the ΔF value for any motor unit pair is limited by the firing rate modulation of the control unit while the test unit is active. Poor rate modulation in the control unit may lead to underestimation of ΔF. To address this possible underestimation, previous work has excluded motor unit pairs in which the rate modulation of the control unit during test unit firing was within 0.5 pps of ΔF (Stephenson and Maluf 2011, Wilson, Thompson et al. 2015).

The present study found removing possibly saturated motor unit pairs had no significant effect on group mean ΔF. This result is consistent with intramuscular findings (Wilson, Thompson et al. 2015). These data suggest that control unit saturation does not have a substantial influence on ΔF value. However, removing possibly saturated pairs, using the method outlined by Stephenson and Maluf (Stephenson and Maluf 2011), may also lead to underestimation of ΔF. Figure 5A shows that the saturated pairs, shown in blue, often have higher ΔF values, due to the mathematical constraints inherent in this method for determining saturation. One possible solution is to calculate the rate modulation in the control unit independently of the ΔF calculation.

### Influence of smoothing method on ΔF

The ΔF calculation relies on smoothed motor unit firing rates. A variety of different smoothing methods have been previously used, and the method chosen to smooth the instantaneous firing rates may influence the ΔF calculation.

While our results show a significant effect of filter type on ΔF value, the range of group mean ΔF across the smoothing methods was negligible (0.2 pps). Filter type also had a significant effect on variance in subject ΔF. Using the fifth order polynomial to smooth instantaneous firing rates provided increased variance in subject ΔF calculation. This is likely due to the fifth-order polynomial method’s increased sensitivity to doublets and erroneous spikes, when compared to hanning or gaussian filters.

Further, the edge effects of these filters play a role in the necessary recruitment and derecruitment time differences between the control and test units. There is a reduced effect of recruitment time difference on ΔF for motor unit pairs that are smoothed using a fifth-order polynomial. Additionally, data smoothed using the shorter 1 s Hanning window reached a plateau in ΔF value at a shorter recruitment time difference than data smoothed using the longer 2 s Hanning and Gaussian windows. These results suggest that a portion of the observed relationship between recruitment time difference and ΔF is due to the smoothing of instantaneous firing rates, in combination with the rapid firing rate acceleration associated with PIC activation. Data smoothed with the fifth-order polynomial were also less sensitive to derecruitment time difference, though to a lesser extent than recruitment time difference. There was minimal difference between smoothing firing rates with the shorter 1 s Hanning window and the 2 s Hanning or Gaussian window. This is possibly due to the slower time course and smaller magnitude of the effect of derecruitment time difference on ΔF.

